# Simplicial and Topological Descriptions of Human Brain Dynamics

**DOI:** 10.1101/2020.09.06.285130

**Authors:** Jacob Billings, Manish Saggar, Jaroslav Hlinka, Shella Keilholz, Giovanni Petri

**Affiliations:** Mathematics and Complex Systems Research Area, ISI Foundation, Turin, Italy; Institute of Computer Science of the Czech Academy of Sciences, Prague, Czech Republic; Department of Psychiatry & Behavioral Sciences, Stanford University School of Medicine, Stanford, USA; Coulter Department of Biomedical Engineering, Emory University and Georgia Institute of Technology, Atlanta, USA; ISI Global Science Foundation, New York, USA

**Keywords:** functional connectivity, time-varying functional connectivity, topological data analysis, persistent homology

## Abstract

Whereas brain imaging tools like functional Magnetic Resonance Imaging (fMRI) afford measurements of whole-brain activity, it remains unclear how best to interpret patterns found amid the data’s apparent self-organization. To clarify how patterns of brain activity support brain function, one might identify metric spaces that optimally distinguish brain states across experimentally defined conditions. Therefore, the present study considers the relative capacities of several metric spaces to disambiguate experimentally defined brain states. One fundamental metric space interprets fMRI data topographically, i.e, as the vector of amplitudes of a multivariate signal, changing with time. Another perspective considers the condition-dependency of the brain’s Functional Connectivity (FC), i.e., the similarity matrix computed across the variables of a multivariate signal. More recently, metric spaces that think of the data topologically, e.g., as an abstract geometric object, have become available. In the abstract, uncertainty prevails regarding the distortions imposed by the mode of measurement upon the object under study. Features that are invariant under continuous deformations, such as rotation and inflation, constitute the features of topological data analysis. While there are strengths and weaknesses of each metric space, we find that metric spaces that track topological features are optimal descriptors of the brain’s experimentally defined states.

**AUTHOR SUMMARY:** Time-Varying Functional Connectivity (TVFC) leverages brain imaging data to interpret brain function as time-varying patterns of coordinating activity among brain regions. While many questions remain regarding the organizing principles through which brain function emerges from multi-regional interactions, advances in the mathematics of Topological Data Analysis (TDA) may provide new insights into the brain’s functional self-organization. One tool from TDA, “persistent homology”, observes the occurrence and persistence of *n*-dimensional holes in a sequence of simplicial complexes extracted from a weighted graph. The occurrence of such holes within the TVFC graph may indicate preferred routes of information flow among brain regions. In the present study, we compare the use of persistence homology versus more traditional metrics at the task of segmenting brain states that differ across experimental conditions. We find that the structures identified by persistence homology more accurately segment the stimuli, more accurately segment high versus low performance levels under common stimuli, and generalize better across volunteers. These findings support the topological interpretation of brain dynamics.

## INTRODUCTION

One of the perennial questions in neuroscience concerns how neuronal signaling generates time-varying experiences. One foundation from which to address this question asserts that brain function emerges from neuronal communication within the context of multiscale neuronal networks. Having access to high-quality whole-brain imaging data, the field of Time-Varying Functional Connectivity (TVFC) (or chronnectomics (Calhoun, Miller, Pearlson, & Adalı, 2014)) offers an empirical approach to characterizing time-varying patterns of mesoscopic neuronal communication (Hansen, Battaglia, Spiegler, Deco, & Jirsa, 2015; Hutchison et al., 2013).

Early computational analysis of brain imaging data observed changes in vectors describing brain **topography** across conditions. FC instead defines a **geometry** among brain regions by computing pairwise similarities from their long-term spontaneous activity measures (Biswal, Zerrin Yetkin, Haughton, & Hyde, 1995). While often the similarity between regions is calculated using the Pearson correlation among spontaneous neuroimaging signals (Biswal et al., 1995; Buckner, 2011; Stoodley, Valera, & Schmahmann, 2010), in general, the idea of brain connectivity can apply to other methods of computing pairwise edges between nodes in the brain. For instance, the present study defines TVFC using instantaneous coherence.

But is the overt geometry of brain imaging data an optimal set of features through which to view and compare brain dynamics? Or, does FC geometry tend to be an idiosyncratic and volunteer-specific descriptor of the brain’s state (Finn et al., 2015)? An alternative perspective observes that an FC **graph** may be treated as a **network**. From here, the analyst may compute graph-theoretic summaries such as centrality, strength, small-worldness, etc. (Bullmore & Sporns, 2009; Farahani, Karwowski, & Lighthall, 2019). However, it is not clear that network properties become clearer when segmenting the brain into more parcels. Rather, the observation of important network properties may require a precise parcellation schema (Gordon et al., 2016).

A more complete picture of neuronal dynamics should account for the brain’s differential establishment, and dissolution, of functionally connected ensembles of brain regions through time. One way to gain this perspective is to consider data as an approximate sampling of an underlying, typically low-dimensional, geometric object, that is, as a **topological space**. In this framework, we may describe the potentially many-body interactions between points or regions of interest using **simplices**. In the simplest and most abstract definition, a *k*-simplex *σ* = [*p*_0_, *p*_1_,…, *p_k_*] is a set of (k+1) points *p_i_* with an ordering. The **topology** of a space is defined by collections of simplices, called **simplicial complexes**, that are closed under intersection (i.e. *X* is a simplicial complex if ∀*σ, σ*′ ∈ *X*, then also *σ* ∩ *σ*′ ∈ *X*). Disconnected holes and cavities are described by the **homology** groups *H_k_* of the simplicial complex: *H*_0_ describes connected components of the complex, *H*_1_ its one dimensional cycles, *H*_2_ three-dimensional cavities, and so on for higher *k*s.

Topological Data Analysis (TDA) attempts to reconstruct the data’s underlying abstract topological space by quantifying the presence and persistence of homological features across different scales (e.g. distances between points, or intensity of correlation between different regions in FC graphs). Such features may include connected regions of a topological space, and its holes in various dimensions, from one-dimensional cycles to higher-dimensional cavities (Battiston et al., 2020; Phinyomark, Ibanez-Marcelo, & Petri, 2017). TDA has been described as “exactly that branch of mathematics which deals with qualitative geometric information” (Carlsson, 2009). In practice, one does not focus on a single complex *X* but rather on a **filtration** 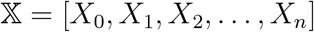, a sequence of nested simplicial complexes, such that *X_i_* ∈ *X*_*i*+1_, which approximates the topological structure at different scales. In this case, the analogues of homological groups are persistent homological groups, which not only capture the presence or absence of a hole, but also at what scale it appears and at what scale—if any—it disappears. In this way, persistent homology generates topological summaries, called persistence diagrams, that can then be used to compute topologically-informed distances between datasets (see Methods).

Re-thinking the more traditional brain dynamics metric spaces from the perspective of topology; values for nodal activity, edge weight, degree strength, etc., are properties that decorate *k*-simplices. Thus, we can consider more traditional metrics as adopting a ‘simplicial approach,’ while a ‘topological approach’ focuses on topological features associated to sequences of simplicial complexes. To compare simplicial and topological spaces of brain dynamics, we leverage pre-existing rest and task fMRI data from 18 volunteers (Gonzalez-Castillo et al., 2015). We compare instantaneous brain images using each of 6 metric spaces—3 simplicial metrics, and 3 topological metrics. Metric spaces are embedded onto 2-dimensions to facilitate statistical tests relating clusters of brain images with common experimental conditions (for more details, see figure 1 and Methods). In part A of figure 2, we report an instance of the embeddings output from the six brain dynamics metrics spaces, that is, the metric space from differential *node* topography, differential *edge* geometry, differential degree *strength*, and also the three topological distances between homology groups in dimensions 1, 2, and 3 (the homology groups *H*_0_, *H*_1_, and *H*_2_). Points often form dense regions associated to certain experimental stimuli. After 256 bootstrap samples of the embedding process, we find that the topological approach excels at distinguishing experimentally distinct brain states.

**Figure 1:**
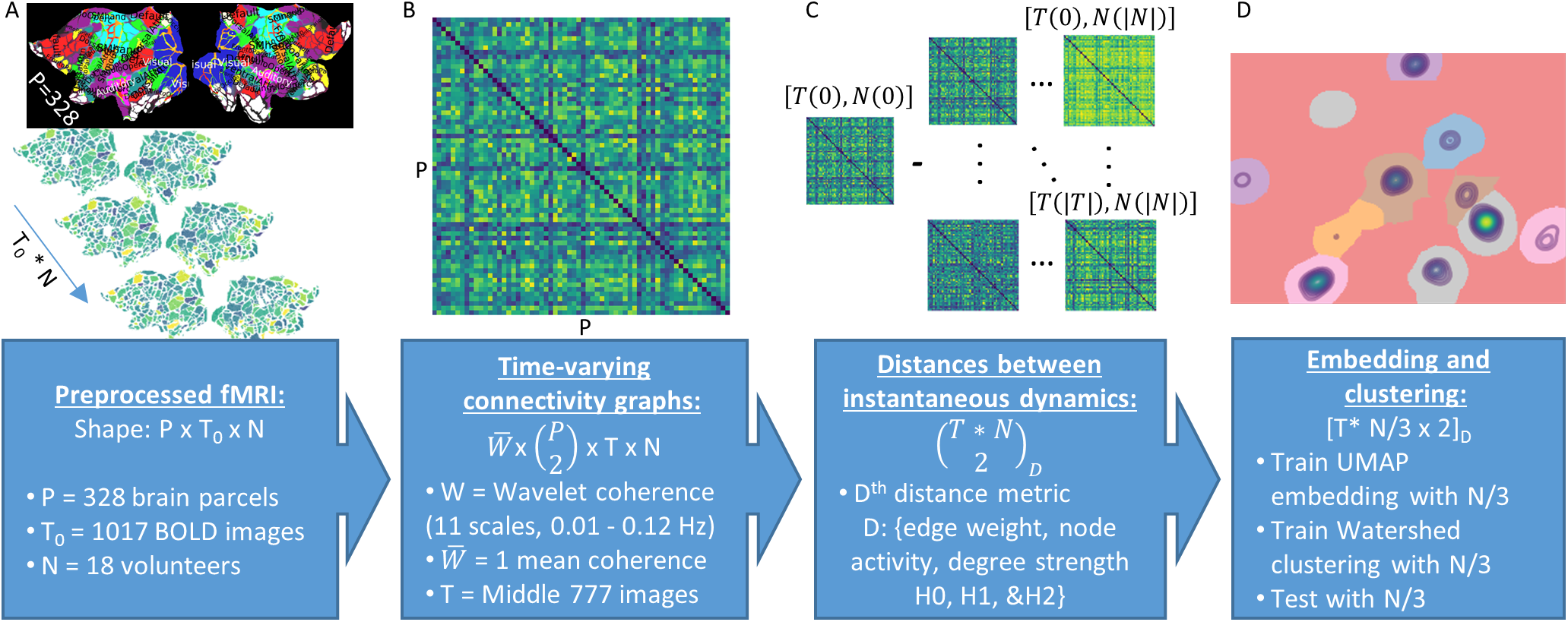
Analysis pipeline. We present the analysis pipeline as a flow diagram in four steps. First, the pipeline accepts preprocessed and spatially segmented BOLD fMRI data as inputs. Then, for each scan, we compute time-varying functional connectivity (TVFC) matrices as the weighted mean of the wavelet coherence between all brain regions, across all time points. Because the wavelet kernel operates over a portion of the time-frequency domain, we remove the outside temporal and spectral edges of the coherence matrix where data padding is required. Next, we compare instantaneous brain dynamics using 6 metrics. Three metrics quantify the similarity among simplex decorations, while the other three compare the lifetimes of persistent homological groups at different dimensions. Finally, we embed each brain dynamics metric space onto 2-dimensions for visualization, clustering, and statistical analysis. To improve seperability among temporally adjacent time points, and to ensure an unbiased clustering of embedded regions, we split volunteers into three groups: 1) an embedding training group, 2) a clustering training group, and 3) a testing group. Statistical results are computed after 256 bootstrapped reinitializations of the volunteer-wise split into the three groups.

**Figure 2:**
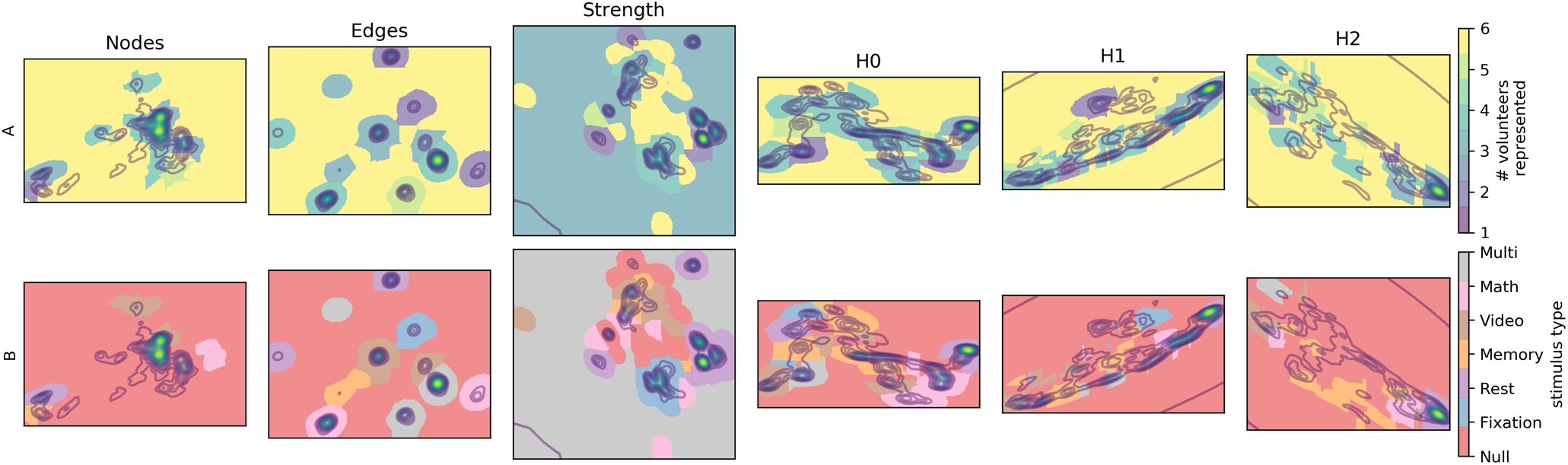
Brain dynamics embeddings for different underlying metrics. We display one realization of the embedded clusters for each of the six metric spaces under investigation. Dense regions of the embedding segment the space into clusters. Clusters are color coded if the underlying points bear statistically significant associations with between 1 to 6 volunteers (part A), or with each of the 5 experimental conditions (part B). (The label ‘multi’ identifies regions independently associated with at least 2 different stimuli).

## RESULTS

### Volunteer-wise representation

As an initial test of the quality of each embedding space, we ask how well the clusters in each embedding generalize across volunteers. To do so, we count the number of points falling into clusters wherein between 1 and 6 volunteers contributed a *not-insignificant* number of points to each cluster. Figure 3 displays the results of this count as percentages with respect to the total number of time points in the test embedding. Following the subsampling and bootstraping schema described in the methods, volunteer-wise generalizability was assessed over 256 independently reinitialized embeddings. Bold lines in figure 3 display the mean, while shaded regions show the 95% confidence interval. A right-skewed distribution indicates increased generalizability, because it means that the densest watershed regions are significantly populated with many volunteers. A left-skewed distribution indicates that most watershed regions are specific to one or few volunteers, i.e., that observed brain dynamics are idiosyncratically related to specific volunteers.

**Figure 3:**
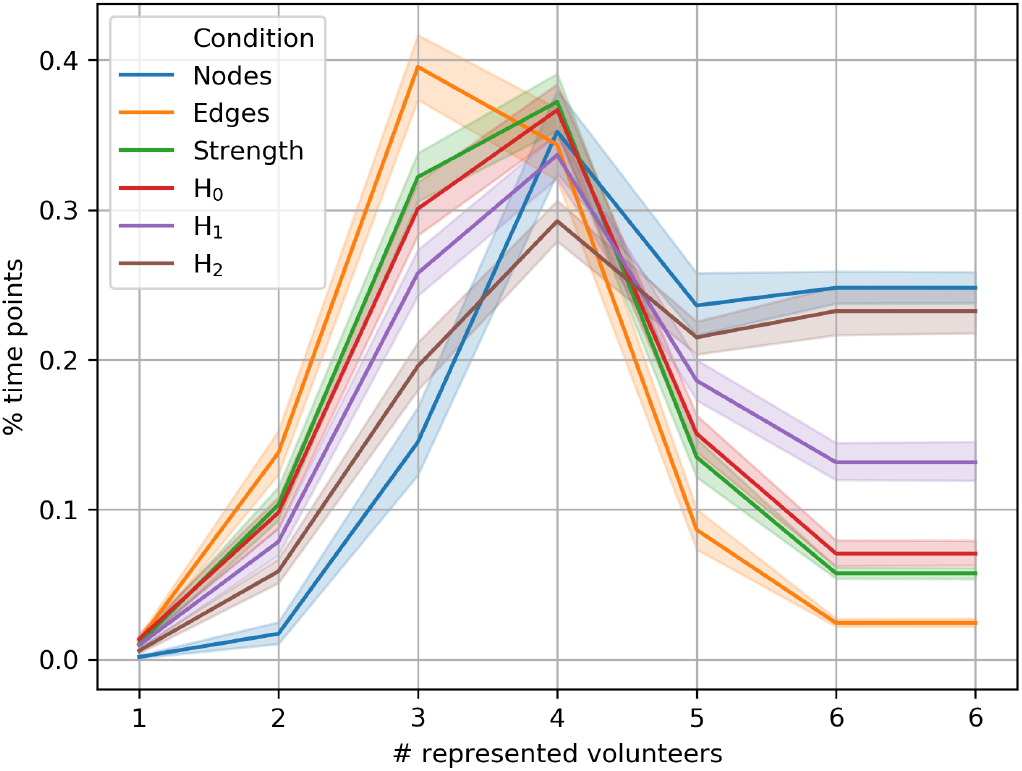
Volunteer specificity of watershed regions. We plot the percentage of time points lying within each of 6 bins. Each bin presents the proportion of points belonging to embedding clusters wherein between 1 and 6 volunteers possessed ‘not an insignificant number of points’ in that embedding cluster (inverse left-tail test). Data are presented as mean and 95% confidence interval over 256 independent samples, each sample from a randomly initialized embedding. Bin 6 is expanded for clarity.

Overall, topological metric spaces offer embeddings that generalize better across volunteers than the other metrics we consider. Not only does homology present right-skewed distributions in figure 3, this category of metrics also aggregates significantly more points into embedding clusters that are general for all 6 volunteers.

It may be possible for metric spaces to generalize too well. For instance, the metric space differing *node* activity agglomerates the largest percentage of time points into bins having between 4 and 6 represented volunteers. However, as will become clear in the next section, this state generalizability comes at the cost of the capacity to distinguish between experimental conditions. Indeed, it appears that the *node* metric space produces embeddings with a single dense core, plus a few distant outliers.

### Stimulus segmentation

A central indicator of embedding quality is the degree to which time points co-localize when belonging to the same stimulus condition. Part B of figure 2 shows an example result of testing watershed clusters against the hypothesis that a significant number of within-cluster points corresponds to any of the 5 experimental conditions. For each stimulus type, figure 4 shows the percentage of points from that stimulus residing in clusters significantly associated with that stimulus (blue boxes). Here again, we report the result as a distribution after 256 independently reinitialized embeddings. Larger percentages of significantly co-localizing points indicate increased capacity to identify brain-states associated with experimental stimuli.

**Figure 4:**
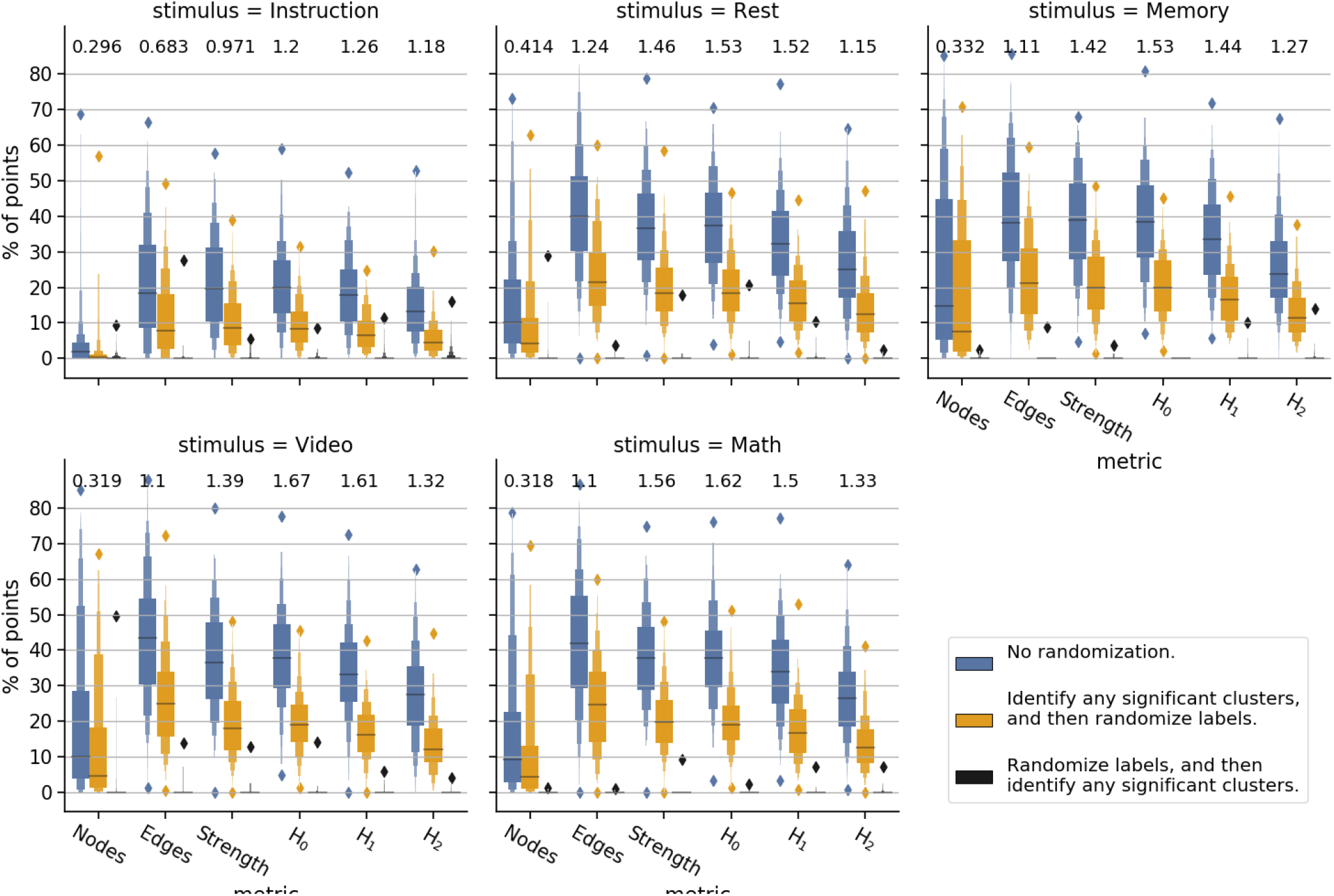
Comparison of task specificity for watershed regions across different metrics. We report the percentage of time points assigned to clusters having a significant amount of points from each experimental condition (blue boxplots). For those same clusters, we report the percentage of points from each experimental condition after randomly permuting point labels (yellow boxplots). Additionally, we report the effect size (Cohen’s D) between these two distributions (values above boxes). A third distribution (black boxplots) shows the false positive rate for identifying significant clusters.

For comparison, we offer two null models computed from randomly permuted point labels. The first null distribution (yellow boxes) permutes point labels among the significant clusters defined previously. It reflects the expected number of points that would randomly collect into the preidentified set of significant clusters. The inclusion of this null model is motivated by the fact that some embeddings clump more points than others into the same watershed region, and would thus hold a larger percentage of points from any experimental condition by default. The effect size (Cohen’s d) between this null distribution and the real distribution provides an indication of how well each embedding isolates brain states induced by distinct experimental stimuli. The second null distribution simply permutes point labels before attempting to find watershed clusters having a significant number of points from any of the 5 experimental conditions (black boxes). This second null distribution provides a good check on the rate of false positives.

Here again, the homology-based embeddings perform very well compared to embeddings constructed from simplicial overlap. This is especially the case for the *H*_0_ metric space which tends to present, over all stimuli, the highest effect sizes. The second highest effect size is found from the *H*_1_ metric space. And the third from the *strength* metric space.

It is interesting to note that, of all the homology-based metrics, the embeddings using Wasserstein distances in *H*_2_ provide the worst segmentation over stimuli. While this may indicate that aspects of TVFC topology are restricted to very low dimensions, the computationally-motivated coarsening of voxelwise information into 328 brain regions also limits the appearance of high-dimensional homologies.

The embeddings over *nodes* produce states that are highly generalizable across volunteers, but that are very poor at distinguishing experimental conditions. In direct contrast, the embeddings over *edges* are the least generalizable across volunteers, but produce embeddings wherein many time points are found in watershed clusters with correctly labeled experimental conditions.

### Task performance

Assuming that differences in performance should be detectable as different brain states under common stimuli, we expect to see large differences between measures of brain dynamics during task time points in which volunteers made fewer or more correct responses. We can test this because the experimental design includes performance metrics for each task, especially the percentage of correct responses for each task block. To do this we computed “mean performance graphs” for each task and each valenced performance level (see Methods). Within each task, performance was valenced as having either more correct responses, or fewer correct responses with respect to a mean split of the performance characteristics for that task from the entire dataset.

Part B of figure 5 displays distances between pairs of mean graphs (across metric spaces and performance levels). Of particular note are the distances computed across the valenced performance levels, but within the same category of metric space (figure 5, white annotations). These values directly measure the sensitivity of each metric space to distinguishing different brain states under common stimuli. Overall, the distance between valenced mean graphs is largest with respect to the topological metric spaces. This is especially true from the perspective of the Jaccard distance (part C of figure 5, lower triangles). From the perspective of the Wasserstein distance in *H*_0_ (upper triangles), the *strength* metric also demonstrates strong cross-valence differences.

**Figure 5:**
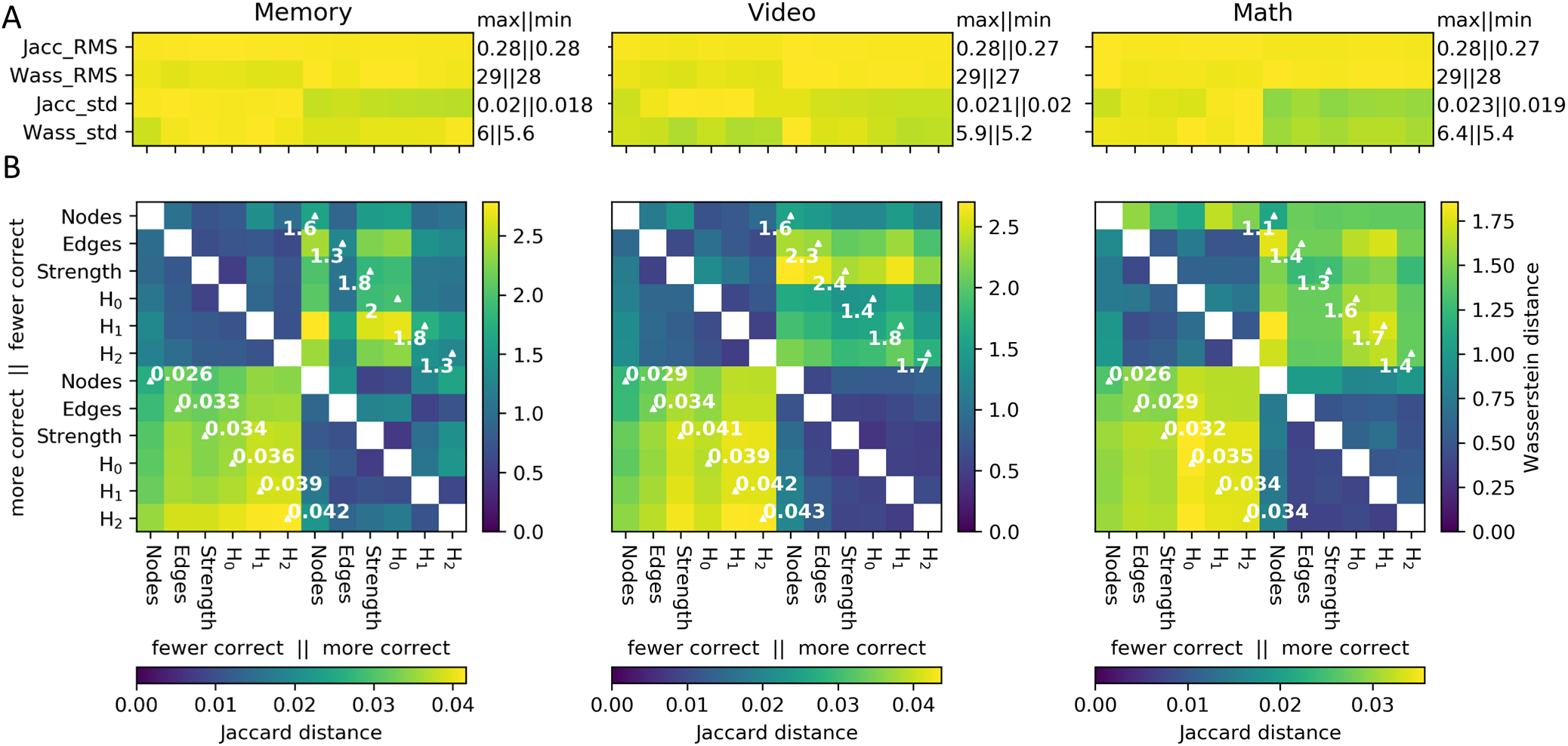
Distances between mean graphs from different performance levels. Mean performance graphs are calculated by taking the mean edge weights for all time points (from any volunteer or condition, and across all embedding reinitializations) located in watershed clusters that are both significantly populated by a given task, and also wherein significantly more or fewer correct responses (with respect to a mean split) were also found for that task (see Methods). Part A of the figure shows the RMS and standard deviations for distances computed between each mean graph versus the set of graphs from which each mean graph was drawn. An annotation is given for the maximum and minimum values in each row. Separate colormaps depict the values in each row. The minimum value is set to 0 for all colormaps. Part B shows distances between the mean performance graphs themselves. Annotations are provided for distances computed within each metric space, but between high performance and low performance mean graphs. For the sake of comparison, distances between mean graphs are calculated with both the weighted Jaccard distance between edges (lower triangle of part B), and also with the sliced-Wasserstein distances between *H*_0_ persistence diagrams (upper triangle). The lower colorbar references the lower triangle, and right colorbar references the upper triangle.

The values in part B of the figure should be compared against summary statistics in part A, and to table 1. Displaying the RMS and standard deviation of the set of distances between each mean graph and their component TVFC graphs provides some indication of the diversity of brain dynamics at times with common stimuli and response characteristics. Compared to table 1, the RMS edge distance between mean graphs and component graphs is below the average *edge* distance across all graphs. By contrast, the RMS Wasserstein distance in *H*_0_ between mean graphs and component graphs approaches the maximum *H*_0_ distance across all graphs. Through the lens of a simplicial approach, mean graphs localize centrally among all graphs. By contrast, through the lense of the Wasserstein distance in *H*_0_, mean graphs are very different from all other graphs. This observation confirms that the simplicial approach and the topological approach are observing very different features of the same datasets.

**Table 1:** Primary statistics, over all distances between pairs of instantaneous brain dynamics.

|  | min | mean | max |
| --- | --- | --- | --- |
| *Nodes* | 5.7 | 38 | 82 |
| *Edges* | 0.0034 | 0.36 | 0.55 |
| *Strength* | 0.0013 | 0.20 | 0.54 |
| $H_0$ | 0.12 | 8.1 | 31 |
| $H_1$ | 0.14 | 2.8 | 9.0 |
| $H_2$ | 0.043 | 1.7 | 6.3 |

### Visualization of homological information

Finally, having identified the high utility of brain-dynamics metric spaces developed from homology to disambiguate group-general brain states, we wanted to gain some insights into what features of TVFC the homology resolves. Owing to the optimal performance of the *H*_0_ metric space, in figure 6, we present a visualization of topological features of a mean performance graph, and also of an instantaneous TVFC graph. Parts A and B of the figure display the *H*_0_ and *H*_1_ homology groups at a single threshold. But the topology of a point cloud should be considered over multiple scales. Part C of the figure gives a sense of the multiscale properties of the lense through topology. Each point in the persistence diagram represents a different collection of homology groups. Interestingly, the observed homology groups in the mean performance graph are shifted to further birth coherence distances than the distrabution of homology groups from the sample TVFC graph. Both distributions of birth and death times are above the threshold for significant wavelet coherence distance, 0.6, as defined relative to an AR1 model of the input data (see part B of supplemental figure 0.1).

**Figure 6:**
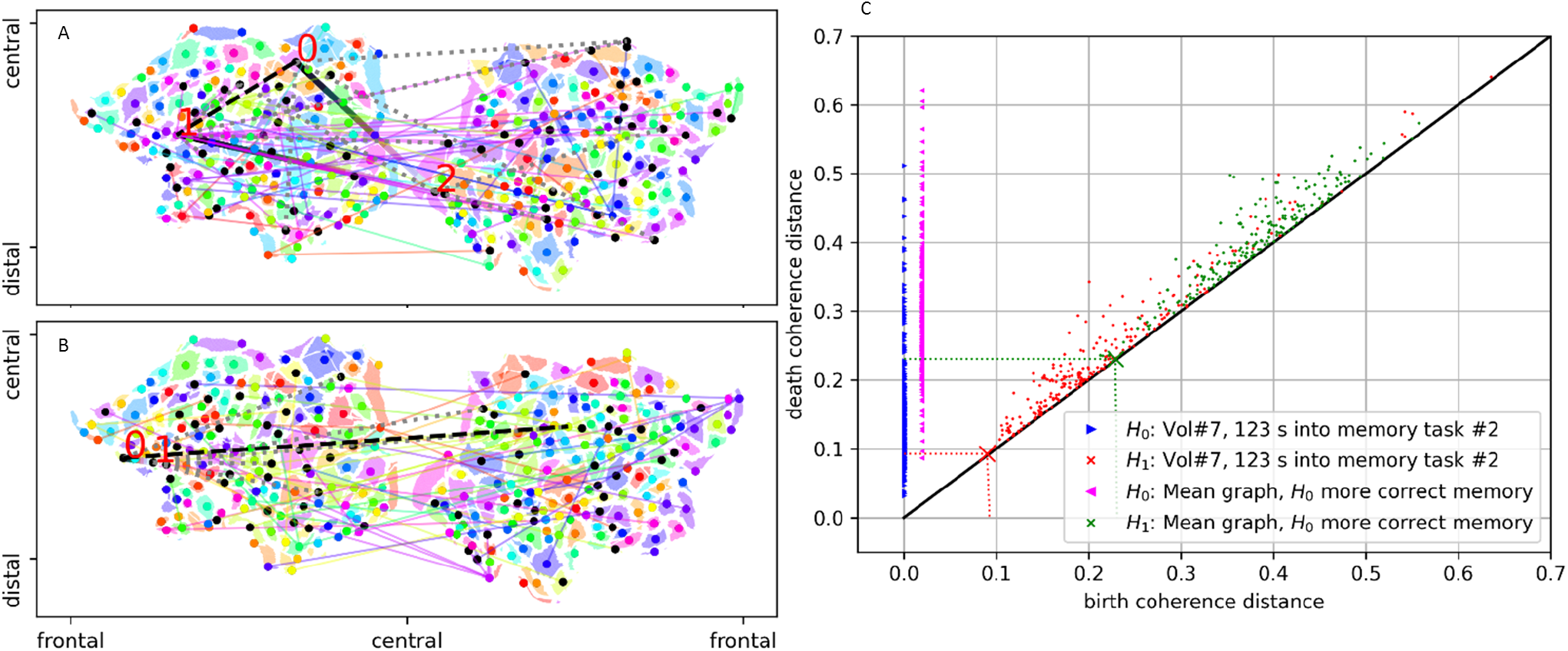
Illustrative examples of persistent homology in *H*_0_ and *H*_1_. Whereas persistent homology operates over a multiscale filtration over inter-node distances, parts A and B of the figure illustrate some of what the algorithm is observing by representing the *H*_0_ and *H*_1_ homology groups at a single scale. The image in part A was computed from the mean graph of more correct memory task responses, as observed by the *H*_0_ metric space. The image in part B represents a single time point consistently identified as a member of the mean graph from part A. The threshold corresponds to the first appearance of a cocycle in *H*_1_. The variegated (‘cubehelix’ colormap) lines in the brain images display the edges involved the cocycle. The red numbers indicate the nodes connected by cocycle edges. Dotted gray lines indicate all edges below this threshold that connect nodes involved in the indicated cocycle. The black dashed line indicates the edge born on or above the threshold that fills in the cocycle. Brain regions are color coded with respect to their clustering with-respect-to an agglomerative clustering with the ‘single’ linkage distance. Light colored lines point between brain regions sharing the same cluster. Colored dots represent the brain region having the largest weighted degree strength of the cluster. Black dots represent the other brain regions of the cluster having less than the maximum weighted degree. For reference, part C of the figure displays the persistence diagrams associated to the graphs from parts A and B. The threshold for the brain images in parts A and B are shown as large ‘x’ markers in part C. The birth time of all *H*_0_ connected components is at zero coherence distance, however, the data are shifted in the ‘x’ axis for clarity.

## DISCUSSION

Brain function is believed to emerge from extensive coordination among brain regions. However, what features typify state-specific brain organization remain a subject of intense and ongoing research (Battaglia & Brovelli, 2020; Lurie et al., 2020). To better understand the correspondence between the methods we use to describe brain dynamics, and the quality of the eventual descriptions, we compared the performance of two broad classes of TVFC metric spaces: one based upon overlap distances between decorated *k*-simplices, and the other based upon *k*-dimensional homological structures. The results of the present study provide evidence that the homology of coherence-based TVFC effectively disambiguates experimentally defined brain states in the population-general brain. By contrast, the performance of approaches based on network and simplicial overlap generally performed worse at distinguishing population-general and experimentally relevant brain states (see figures 3 and 4).

### Intrinsic geometries

Given a good space for representing brain dynamics, it is possible to observe stereotypical brain states between more subtly different conditions. Utilizing the same dataset as the present study, Saggar et al. (2018) computed a distance between node activities to visualize two-dimensional mappings of within-volunteer temporal similarity. In the majority of cases, the visualization depicts even transitions across time points. Smooth transitions over short distances are clearly depicted during the resting state. Smooth transitions are also a feature of most temporally adjacent transitions during task states. However, for some volunteers, the mapping depicts modularized transitions within the context of a single experiment.

Using a complimentary dataset, Billings et al. (2017) also computed maps of node activity distances. Distances were mapped across a population of volunteers. Even at the group level, a general trend was observed of variable activity punctuated by moments of clear transitions between focal brain states. Similarly, a sample of the *nodes* embedding shown in figure 2 contains *O*10 very dense nodes distributed along the circumference of a more sparsely populated embedding.

### Towards a topological view

While studies implementing simplicial metrics evidence that brains select conserved dynamical patterns towards the production of brain function, the empirical and theoretical support for emphasizing homological and other topological descriptors has prompted several authors to reinterpret neuronal dynamics from a topological perspective (Curto, 2017; Giusti, Ghrist, & Bassett, 2016; Lerda, 2016; Rasetti, 2017; Reimann et al., 2017; A. E. Sizemore, Phillips-Cremins, Ghrist, & Bassett, 2019; Stolz, 2014). A. E. Sizemore et al. (2018) evidence that **cliques** and homological cavities in the mesoscopic space of structural brain images reflect known brain networks. Further evidence that cliques and homologies encode microscopic interactions among neuronal circuits have been discovered within the hippocampal place field (Basso, Arai, & Dabaghian, 2016; Dabaghian, Brandt, & Frank, 2014; Giusti, Pastalkova, Curto, & Itskov, 2015) and in the somatomotor representation of the head (Chaudhuri, Gerçek, Pandey, Peyrache, & Fiete, 2019). The present results provide further support for the utility of the topological approach to discern the evolution of brain states through time, thus to possibly improve our comprehension of the brain’s multiscale self-organization.

As a quantitative tool, persistent homology is tailor-made for defining topological similarities among metric spaces (Carlsson, 2009). Indeed, fMRI studies have implemented persistent homology to discern group-level FC differences in task performance (Ibáñez-Marcelo, Campioni, Phinyomark, Petri, & Santarcangelo, 2019), and with respect to pharmacological treatments (Petri et al., 2014a). Similar findings are observed in MEG data (Duman et al., 2019). Stateful segmentation was also achieved from homological features in *H*_0_ for 8-channel EEG TVFC as volunteers engaged in a visuo-motor task (Yoo, Kim, Ahn, & Ye, 2016).

### Visualizing topology

Certainly, functional connectivity describes a multiscale process. And while there are ongoing questions regarding the pathways through which otherwise structurally distributed brain networks form TVFC networks (Damoiseaux & Greicius, 2009); the development of data-drivin functions that operate over spectral and spatial features of complex networks may drive new insights. The view from homology may be especially useful when topological features are expected to be important, that is, when one expects multiple scales of patterned connectivity among clusters in *H*_0_, and/or higher order (dis)-connected cycles in *H*_1_ and above. In pursuit of this hypothesis, it may be useful to start with a more dense spatial sampling over brain regions. Also, the expansion of the spectral data into a multi-layer graph may improve stateful representations. In any case, the present observation of meaningful homology in *H*_0_ may relate to the fundamental description of brains as functioning through multiple scales of interacting brain regions. Given the theoretical significance of homology in *H*_0_ (e.g., multiscale clustering), and it’s computational speed increases relative to computing homology in *H*_1_ and above; it appears to be worthwhile to use persistent homology in *H*_0_ as a general tool for describing and comparing brain states.

### Limitations and future directions

Future research should strive to make a more detailed catalogue of the homologies that commonly appear among brain regions. While the present study resorted to a very coarse brain parcellation to visualize homology (see figure 6), it is not clear how stable these minimal cycles are. Indeed, it is not clear that 333 parcels provides a maximal resolution of brain dynamics. In theory, more parcels should enhance the capacity for persistent homology to distinguish brain states; albeit, up to some plateau. By contrast, elementwise operations over simplicial decorations benefit from clustering (Glasser et al., 2016; Gordon et al., 2016) and unmixing (Kunert-Graf et al., 2019; Smith et al., 2009). Future should utilize this stability property of TDA to catalogue the stability of cycles across multiple scales of parcellation. Another limitation of the present study is the reliance on clustering in the low-dimensional space. Even while low-dimensional embeddings provide an efficient means for visualizing data, there is always some loss of information. For instance, the UMAP method for embedding point cloud data transduces an explicit nearest-neighbor approximation of the high-dimensional simplicial complex into the low-dimensional space. This approximation may be causally related to the observation that metric spaces based upon, especially, 1-dimensional simplicial overlap organize into temporally-adjacent clusters. While edge overlap may be a volunteer-specific trait. And while the trait may be partially alleviated by deconvolution of the volunteer-specific hemodynamic response function, future work that biases the low-dimensional embedding in a more appropriate way—perhaps by learning a transductive vector embedding as in Bai et al. (2019)—may offer some additional improvements. In any case, approaches that circumvent dimensionality reduction entirely by operating in the native high-dimensional space may offer the most general solution to the loss of information during low-dimensional embedding.

Finally, it is always interesting to consider more concise multispectral decompositions than provided by Morlet wavelet kernels. Perhaps kernels that imitate the canonical hemodynamic response function would offer a more compact representation of fMRI data. Also, while the Morlet wavelet is roughly symmetric, it may be useful to implement asymmetric filters that place more emphasis on information from more recent time points.

### In conclusion

To understand the dynamic self-organization of complex systems like the brain, it helps to view system dynamics through lenses that highlight the presence and the structure of complexes. Given the kinds of weighted graphs typical of TVFC analysis, persistent homology is well-suited for interpreting complexes of brain regions. The view from homology outperforms more traditional graph metrics —like the activity measures of 0-dimensional nodes, and like the weights of 1-dimensional edges—at the task of segmenting experimentally defined brain states into patterns that generalize well across multiple volunteers. The utility of these data-drivin multiscalar methods inspires additional research into the topology of high-dimensional connected objects.

## METHODS

As described in figure 1, our procedure unfolds across 4 steps:

1. Acquire task and resting-state BOLD fMRI data from a group. Apply minimal preprocessing.
2. Compute TVFC as instantaneous coherence.
3. Differentiate instantaneous brain dynamics via each of 6 metrics:

a. Euclidean distance between *node* topographies
b. Weighted Jaccard distance between *edge* geometries
c. Weighted Jaccard distance between the weighted degree *strength* of networks
d. Sliced-Wasserstein distance between topographic persistence diagrams in *H*_0_
e. Sliced-Wasserstein distance between topographic persistence diagrams in *H*_1_
f. Sliced-Wasserstein distance between topographic persistence diagrams in *H*_2_
4. Embed distance onto 2-dimensions for visualization and statistical analysis

### Data acquisition and preprocessing

To discern the relative capacities of a range of distance metrics to disambiguate the dynamical brain-states induced by stimuli, for the present study, we adopted a dataset acquired during the presentation of multiple experimentally defined tasks. The present study benefited from scans acquired continuously over relatively long time spans as the process of spectral filtration requires complete overlap between the signal and the filtration kernel to avoid affects at the undefined edges of the time series. And, whereas we are interested in signals in the low-frequency fluctuation range (1/100 seconds^2^), we required scans to be at least longer than 200 seconds.

The data acquired by Gonzalez-Castillo et al. (2015) meet these criteria. These data have been publicized as an open-access dataset through the XNAT neuroimaging database (https://central.xnat.org; Project ID: FCStateClassif). Here, we briefly summarize the dataset as follows: 18 volunteers were scanned continuously over 25.5 minutes (7 Tesla, 32-element coil, gre-EPI, TR=1.5s, TE=25ms, 2mm isotropic). Preprocessing was performed to transform individual datasets into a common MNI space and to remove artifacts from slice timing, motion, linear trends, quadratic trends, white matter signals, and csf signals. Data were spatially smoothed using a 4mm FWHM Gaussian filter. They were temporally band-pass filtered to between 0.009 Hz and 0.08 Hz. Finally, images were downsampled to 3mm isotropic, and normalized to common (MNI) coordinates. Data were acquired in compliance with a protocol approved by the Institutional Review Board of the National Institute of Mental Health in Bethesda, MD. For complete preprocessing details, please refer to Saggar et al. (2018). In addition to the aforementioned steps, voxelwise data were spatially aggregated onto an atlas of 333 brain regions (Gordon et al., 2016). Up to 5 brain regions contained no information from some volunteers, and were excluded from all datasets for the remainder of the analysis. (Numbers 133, 296, 299, 302, and 304, indexed from 0. See also the missing patches in figure 1, part A) Thus, the finest granularity of study results are over 333-5=328 brain regions. During the scan, volunteers interacted with 3 block-design tasks and one rest stimulus. Each task was presented twice. Each task presentation lasted 3 min, and was proceeded by a 12s instruction block. Tasks included: ‘video,’ watching videos of a fish tank while responding to a visual target; ‘math,’ computing algebra problems; and ‘memory,’ a 2-back memory task with abstract shapes. A ‘rest’ stimulus was also included, and entailed the presentation of a fixation cross for 3 minutes. Stimuli were randomly ordered in a fixed sequence for all volunteers. For each task block, performance metrics were collected, including the percentage of correct responses.

### Time-varying connectivity

Considering that individual frequency bands develop significantly different FC parcellations (Billings et al., 2018) and different connectivity hubs (Thompson & Fransson, 2015). And, considering that neuroelectric activity is intrinsically rate coded. The delayed and (hemodynamic response function) band-pass filtered version of neuroelectric activity that is the BOLD signal is likely to retain some rate-coded information. Given these observations, the present study recasts the BOLD signal from each brain parcel in terms of time-frequency spectrograms generated through the use of the Continuous Wavelet Transform (CWT)

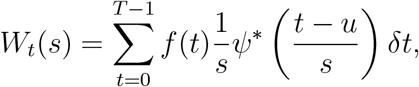

where ·* indicates the complex conjugate. By adjusting the time localization parameter *u* and the scale parameter *s* for the wavelet kernel *ψ*, the CWT affects a multiscale decomposition of input signal *f*(*t*) for all times *t* ∈ *T*. For the present study, the filterbank comprised 15 scales log-distributed between 0.007 and 0.15 Hz.

Following Torrence, Compo, Torrence, and Compo (1998), symmetric wavelets will produce similar coherence values. And without strong support for any particular wavelet kernel, we adopt the complex Morlet wavelet as the CWT kernel. The filter is a plane wave modified by a Gaussian, *ψ* = *e*^*iω*_0_*t*/*s*^*e*^−*t*^2^^/(2*s*^2^). And we set the base frequency to *ω*_0_ = 6. Following Farge and Marie (1992), an *ω*_0_ > 6 ensures the function’s non-zero average is outside machine precision (Farge & Marie, 1992). Spectral selectivity increases with increasing *ω*, at the expense of decreased temporal selectivity (e.g., sharper filters require more temporal support). Thus, a base frequency of *ω*_0_ = 6 ensures maximal temporal resolution.

A complex valued kernel computes instantaneous amplitude and phase information. From there, it is possible to compute wavelet coherence as follows. For a pair of complex-valued spectrograms, *W^X^* and *W^Y^*, the quantity 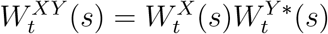 is the cross-wavelet spectrum. Its absolute value, 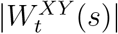, is the cross wavelet power which represents the shared power between signals at scale *s* and time *t*. Coordinated changes in amplitude may be computed in terms of the wavelet squared coherence,

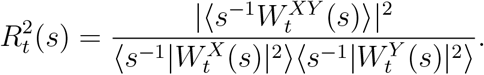

The functional 〈·〉 indicates smoothing in both time and scale. The factor *s*^−1^ is used to convert to scale-dependent energy densities. The wavelet squared coherence is an instantaneous and multispectral analogue of the Pearson correlation (Marwan, Thiel, & Nowaczyk, 2002; Torrence et al., 1998; Torrence, Webster, Torrence, & Webster, 1999). Its values range between 0 (completely incoherent) and 1 (completely coherent). While it is theoretical possible to treat TVFC as a multilayer graph having as many layers as spectral scales, practical computational concerns prompt us to concatenate multispectral coherence into a single broadband average. To do so, we take the weighted mean of the wavelet squared coherence with respect to the normalized cross wavelet power:

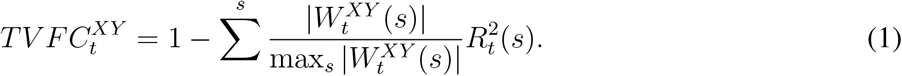

Normalizing the cross wavelet power ensures that the mean coherence remains bounded between 0 and 1. The peak of the mean cross wavelet power occurs in the frequency range between 0.01 and 0.02 Hz and (see part A of supplemental figure 0.1). TVFC graph edges are 1 minus the power-weighted coherence to represent coherence distances between brain regions.

To account for the cone of influence at the temporal edges of the wavelet filtration, as well as the loss of precision at the temporal and spectral edges of the smoothed coherence data, the outside 120 time points and the outside 2 scales are dropped before taking the summation in equation 1. The removed time points include one whole “rest” block, and one whole “video” block. Coherence graphs are thus available for the middle 777 images of the scan, and for 11 spectral scales between 0.0095 and 0.1 Hz.

### Distance metrics comparing brain dynamics

#### Theory

Having constructed TVFC graphs for all included time points and for all volunteers, we pursue two broad alternatives for comparing brain dynamics. The first is related to elementwise differences between the decorations (e.g. weights) applied to graphs. And the second relates to common topological features. To describe in detail these two views, it is useful to supply some definitions.

A graph *G* = (*V, E*) represents a set of *V* nodes interconnected by *E* edges. Nodes and edges may be decorated with properties such as value, weight, directionality, sign, layer, degree centrality, degree strength etc. A collection of *k* completely interconnected nodes forms a clique, *C*. In the following, we identify cliques with geometric primitives called ‘simplices’ in standard fashion (Petri et al., 2014b; Petri, Scolamiero, Donato, & Vaccarino, 2013); that is, to a clique of *k* + 1 nodes we associate the corresponding *k*-simplex, *σ_k_*. For instance, 2 connected nodes form a 2-clique. The surface enclosing a 2-clique is a 1-simplex, i.e., an ‘edge’. A 2-simplex formed by a clique of 3 connected nodes is a ‘filled triangle’, and so forth for higher-order simplices.

Formally, a simplicial complex is a topological space, 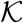, composed of all *σ_k_* and their subfaces. Along the same lines, a clique complex, *Cl*(*G*), is a simplicial complex formed from an unweighted graph *G* by promoting every *k*-clique into a (*k* − 1)-simplex. Holes in dimension *k* may develop within the boundaries established by closed chains of (*k* − 1)-simplices. Such holes are called ‘homologies.’

The ‘topological approach,’ TDA, includes methods for identifying topological features of an abstract geometric object represented by a data sampling. By contrast, the more traditional approach to comparing brain dynamics constitutes a ‘simplicial approach’ that directly compares the decorations applied to sets of simplices.

#### Homology

The boundary of a homology is termed a, ‘homological cycle’ or ‘generator.’ To illustrate the concept, consider the case of four nodes connected in a cycle such that each node has exactly two edges. The nodes form neither a 4-clique nor a 3-dimensional simplex because there are two missing edges. Rather, these nodes form a connected cycle that is the boundary of a 2-dimensional hole. This void space is also called a *homology* in dimension 1 (i.e., formed by a set of 1-d edges). The *k^th^* homology group, 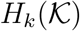, describes the (*k* + 1)-dimensional holes bounded by chains of *k*-simplices. For example, the *H*_1_ homology group are the holes bounded by edges in 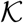; *H*_2_ are the voids bounded by filled triangles; etc.

The term ‘homology’ follows from the Greek ‘homo,’ the same, and ‘logos,’ relation, to indicate that the hole belongs to an equivalence class that is categorically the same across continuous deformations that neither break the boundary nor create new simplices spanning the boundary: e.g., inflation, compression, rotation, and translation. Different representative cycles may therefore exist that describe the same homological cycle. For instance, a very elastic coffee cup could be continuously contracted into the shape of a donut, as they share the same toroidal topology. For the sake of convenience, a homological cycle is often represented as the minimal representative cycle (Guerra, De Gregorio, Fugacci, Petri, & Vaccarino, 2020; Petri et al., 2014b).

#### Simplicial distances

The first approach, which we will denote as ‘simplicial,’ computes an average of the elementwise differences between the decorations applied to each *k*-simplex in the complex. For example, in the present study, we compute the weighted Jaccard overlap distance between the weights of TVFC *edges* as

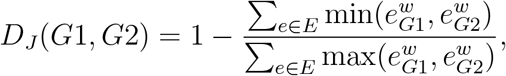

where 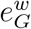 is the weight of the *e^th^* edge in graph *G*.

Further we compute distances between the explicit 0-dimensional values decorating each node; e.g. with respect to the signal activity of each *node*. Specifically, for each point in time, we treat the absolute values of multispectral wavelet coefficients from all brain regions as an ordered vector. We then compute the Euclidean distance between vectors from different points in time.

The third distance is inspired by previous work on relations between graph networks and homological cycles. Lord et al. (2016) demonstrate that the nodes’ weighted degree (also called *strength*) is significantly correlated with the frequency and the intensity with which nodes participate in the shortest representatives of homological cycles. The third distance is thus the weighted Jaccard distance between vectors of the node-wise weighted degree, also called the *strength*, of each TVFC graph.

#### Homological distances

While many TVFC studies regard only the graph’s connectivity as the feature of primary import, TDA provides a suite of tools to further develop network properties into conserved higher-order structures in point-cloud data (Carlsson, 2009; Edelsbrunner, Letscher, & Zomorodian, 2002; Patania, Vaccarino, & Petri, 2017) and in weighted networks (Chung, Lee, Di Christofano, Ombao, & Solo, 2019; Petri et al., 2013; A. Sizemore, Giusti, & Bassett, 2017).

Homology is defined on simplicial complexes. In the case of persistent homology of weighted graphs, simplices are added to the complex incrementally, and appear at and beyond some threshold. Varying this threshold allows to track how homological features appear and persist across thresholds (Petri et al., 2013). A complete representation of homolocial features within some range of thresholds is called a ‘filtration.’ By observing topological features over a filtration, “persistent homology” allows to take a multiscale view of the data which accounts for both the explicit connectivity structure of the system, as well as the relative importance of ensembles of connections that emerge over some range of scales.

Formally, we define the Vietoris-Rips simplicial complex 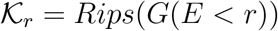 as the clique-complex of the weighted graph *G* composed after removing all edges, *E*, longer than *r*. From this, we may recover the complex’s *k*-dimensional homology group, 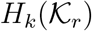. Within the boundaries of thresholds *a* and *b*, let [*r_a_*,…, *r* – *ϵ, r*,…, *r_b_*] be the longest series wherein any 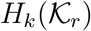 and 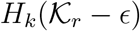 are not identical. The ordered set 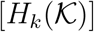 defines a ‘filtration’ over *G*. A homology class *α* ∈ *H_k_* is said to be *born* at radius *u* if a class of homotopy equivelant homologies is not supported in 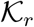 for any *r* < *u*. The homology class *α* is said to *die* going into 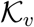 if *υ* is the lowest index wherein at least one (*k* + 1) – *clique* is established within the boundary of the homology. Persistent homology was computed using version 0.4.1 of the Ripser package as bundled with the Scikit-TDA toolbox for python (Tralie, Saul, & Bar-On, 2018). Ripser finds it is faster to compute cohomology, the covariant **functor** of homology. Thus the algorithm computes cocycles in *H_k_* that track the disappearance of *σ*_*k*+1_ along the reversed filtration De Silva, Morozov, and Vejdemo-Johansson (2011).

The persistent homology of a filtration over *G* is summarized by collecting the birth/death pairs of *k*-dimensional homology classes as points (*u, υ*) in a “persistence diagram”. It is naturally possible to compute a persistence diagram for each simplicial dimension up to the maximum dimension of the simplicial complex. But because the computational load to calculate persistence homology increases exponentially with the homology dimension, we limit the present study to the investigation of persistence homology in dimensions 0, 1, and 2. The case of 0-dimensional persistence diagrams—corresponding to 0-dimensional holes, that is, disjoint sets of connected nodes—is particularly interesting as the homological classes are slices through an agglomerative clustering among nodes when using the ‘simple’ linkage distance.

Persistence diagrams can, themselves, be endowed with a metric structure. This means that it is possible to measure distances between persistence diagrams. Such distances encode how different the homological structures of two TVFC graphs are. One such distance is a multi-dimensional analogue of the earth-mover distance, known as the sliced-Wasserstein distance (Carrì, Cuturi, & Oudot, n.d.). The sliced-Wasserstein distance between persistence diagrams is bounded from above by the total distance between the associated topological spaces (Mileyko, Mukherjee, & Harer, 2011). In the present study, for each pair of persistence diagrams of a given dimension, we calculate the average Wasserstein distance, over 20 slices (see Carrì et al. (n.d.) for details). That is, for all pairs *G^i^* = *G^j^* we compute, 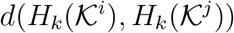.

### Visualization/Output

Having developed metric spaces to compare simplicial and homological brain dynamics, we want to assess their relative capacities to represent apparent brain states. To this end, we embed each metric space onto a 2-dimensional manifold using the Uniform Manifold Approximation and Projection (UMAP) algorithm (McInnes, Healy, & Melville, 2020). As illustrated in figure 2, the embedding process facilitates state-space visualization and segmentation. UMAP approximates a metric space’s *n*-dimensional manifold in three steps. First, the algorithm calculates the k-nearest neighbors of each point. Second, each neighborhood is promoted to a local simplicial complex. Third, the algorithm searches for the *n*-dimensional distribution of points that best approximates the original simplicial complex. This search is conducted over successive iterations, with the initial position of low-dimensional points derived from a random distribution.

To better understand the distribution of points in the resulting embedding spaces, we transformed point clouds into a Gaussian distribution and estimated clusters via a watershed transform. An illustration of watershed clustering is found in part B of figure 2. The Gaussian grid size was initially set to 256×256. The number of grid points in the dimension having the smaller range was trimmed to maintain the aspect ratio of the embedding. The Gaussian kernel bandwidth factor was set to 0.08. The watershed transform marks the local densities as cluster centers, then grows clusters by adding adjacent pixels whose directed gradient is maximal in the direction of the cluster center.

### Subsampling and Bootstrapping

In the present study, we were concerned with resolving 2-dimensional embeddings that generalize across volunteers, while also segmenting experimental stimuli. One challenge in the way of resolving this ideal embedding is that brain states tend to change slowly through time. An example of this issue is shown in supplemental figure 0.2 for the metric between TVFC *edges*. Temporal similarities draw the distance between adjacent time points closer than the distance between two different volunteers experiencing the same stimuli. For dimensionality-reduction algorithms like UMAP and tSNE that leverage nearest-neighbor approximations, the attractive force between temporally-adjacent time points can force the embedding to over-emphasize information about the order of the scanning sessions when attempting to resolve population-wise brain states.

To help disentangle graphs representing intrinsically similar brain states from those that are simply autocorrelated, we subsampled our dataset in several ways. Statistics over the results could then be generated via bootstrapping, with 256 random permutations of data subsamplings.

Volunteer-wise scans were split into three equal groups. The first group supplied data to train the UMAP embedding. The second group supplied data to segment the space of the embedding into watershed clusters. The third group supplied data to test how metric spaces segment brain states during contrasting experimental conditions.

The data were also split in time. To balance the number of time points from each experimental condition, one of each of the repeated mathematics and memory tasks were removed, at random, from each volunteer’s dataset. Also, embeddings were trained using 100 time points from the remaining 537 time points of 6 volunteers. These 100 training points were selected to emphasise maximal temporal separation.

### Statistical Analysis

Watershed clusters provide a data-driven basis for hypothesis testing over the likelihood that certain metadata labels—that is, volunteer number, stimulus type, and performance—were more or less likely to be found in a given embedding region. For all statistical tests, we generated null distributions by randomly permuting the labels of cluster points (e.g. volunteer number, experimental condition, etc.) 300 times. This procedure obtained a mean and standard deviation that indicates the labels we should expect to find by chance in any given cluster. The significance threshold was always set to an *α* = 0.05. Bonferroni correction was applied relative to the number of simultaneous tests performed. And the total number of clusters was *O*100 in each embedding.

Tests related to volunteer co-localization calculated significant volunteer-wise under-representation in each cluster (left-tail test, Bonferroni correction equal to the number of volunteers (6) times the number of clusters per embedding (*O*100)). Tests related to stimulus co-localization identified clusters that were more than likely to contain time periods during each stimulus condition (right-tail test, Bonferroni correction equal to the number of stimulus conditions (5) times the number of clusters in each embedding (*O*100)). Tests related to task performance were conducted for each task condition independently, and were confined only to the clusters that were significantly more likely to contain points from the task being tested (two-tailed test, null-distribution is the mean and standard deviation of task performance, Bonferroni correction equal to the number of clusters showing significantly many within-condition time points (*O*10)).

### Secondary Statistics over Mean Graphs

It is possible to generate mean FC matrices from select time points of TVFC graphs. For instance, the mean TVFC graph over all time points reveals the average coherence between regions. Condition-dependent mean graphs such as that over all rest conditions may also be calculated. In the present study, we were particularly interested in mean graphs calculated with respect to within-task performance levels.

Given the identification of clusters significantly associated with task performance. For each task, and for each cluster associated to the task, we tested whether the task-specific points within that cluster contained significantly more or fewer correct responses than the mean percentage of correct response for all of that task’s time points (no Bonferroni correction). For each task, every time point from clusters having significantly more correct responses is stored into a task-specific list. The same process occurs for clusters showing fewer correct responses. The mean TVFC graph from each list constitutes a ‘mean performance graph.’ Mean performance graphs may be compared to one another to measure a difference between apparent brain states.

## ACKNOWLEDGMENTS

This work could not have proceeded without the insights from Dr. Alessio Medda on wavelet theory, nor without the essential works of our global communities.

## TECHNICAL TERMS

**topography** The vector if a multivariate signal measuring a system at a given instant.

**geometry** The study of distance functions.

**graph** A finite set of nodes, equipped with a finite set of edges.

**network** A graph where-in edges convey the property “interacts with.”

**topology** A collection of subsets of a set.

**topological space** A totality of two elements: a set of points, and a topology on this set.

**clique** A set of *k* nodes.

**simplex** The *k*-dimensional convex hull of a clique of *k* + 1 nodes.

**simplicial complex** A collection of multiple simplices.

**homology** A *k*-dimensional hole bounded by cyclically connected (*k* + 1)-dimensional simplices.

**filtration** Varying the threshold parameter of a weighted graph to resolve simplicial complexes with altered homology.

**functor** A function between categories which maps objects to objects and morphisms to morphisms. Functors exist in both covariant and contravariant types.

## SUPPORTING INFORMATION

**Figure 0.1:**
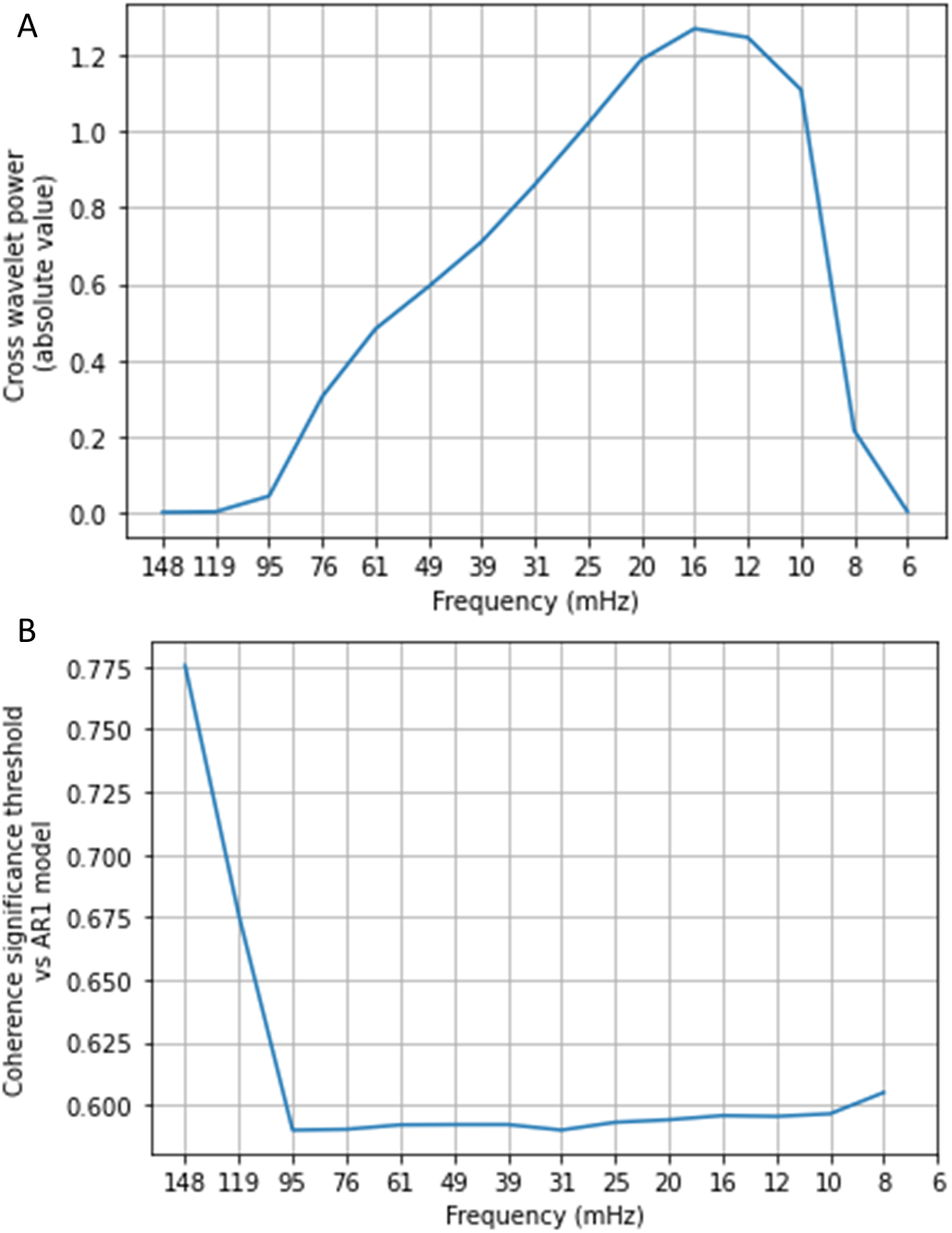
Describes the mean values of the input data across frequency bands. Part A of the figure displays the absolute value of the mean cross wavelet power. Frequencies between 0.01 and 0.02 Hz are most likely to contribute to the mean wavelet coherence used to quantify TVFC edges. Part B of the figure displays a significance threshold level indicating significantly high coherence. The threshold was calculated as an average against the background power spectrum. A distribution over the background power spectrum was calculated from 300 lag-1 approximations of each time series. Following Torrence et al. (1998), the 95% confidence interval is the product of the background power spectrum and the 95th percentile value of a chi-squared distribution with two degrees of freedom.

**Figure 0.2:**
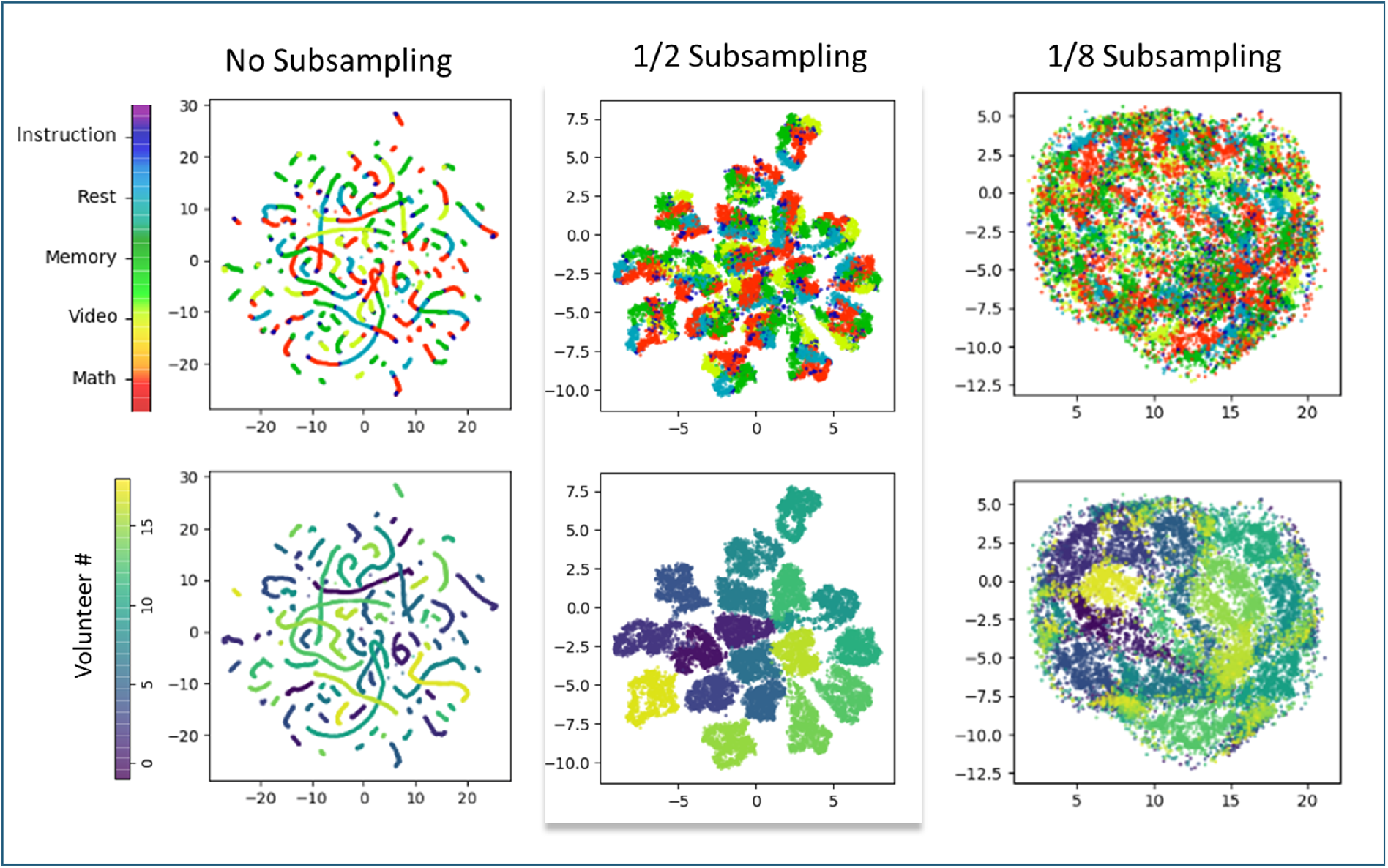
Observes an embedding of the weighted Jaccard distance between edges at several levels of subsampling. With no subsampling, successive time points from within the same scan retain strong attractive forces in the UMAP nearest-neighbor embedding. This produces string-like masses. Subsampling the data by half retains the similarities present within each volunteer’s scan, thereby grouping volunteers into their own cluster. A noticeable degree of volunteerwise clustering—e.g., temporal selfsimilarity—is still present when seeding the embedding with 1/8 the total number of data points. None-the-less, some mutual attraction does emerge among edge-centric brain states measured across multiple volunteers.

## REFERENCES

Bai, Y., Ding, H., Qiao, Y., Marinovic, A., Gu, K., Chen, T.,… Wang, W. (2019). Unsupervised Inductive Graph-Level Representation Learning via Graph-Graph Proximity.

Basso, E., Arai, M., & Dabaghian, Y. (2016). Gamma Synchronization Influences Map Formation Time in a Topological Model of Spatial Learning. PLOS Computational Biology, 12(9), e1005114. Retrieved from https://dx.plos.org/10.1371/journal.pcbi.1005114 doi:10.1371/journal.pcbi.1005114

Battaglia, D., & Brovelli, A. (2020). Functional connectivity and neuronal dynamics: insights from computational methods (Tech. Rep.). Retrieved from https://hal.archives-ouvertes.fr/hal-02304918

Battiston, F., Cencetti, G., Iacopini, I., Latora, V., Lucas, M., Patania, A., … Petri, G. (2020). Networks beyond pairwise interactions: structure and dynamics. Physics Reports.

Billings, J., Medda, A., Shakil, S., Shen, X., Kashyap, A., Chen, S., … Keilholz, S. D. (2017). Instantaneous brain dynamics mapped to a continuous state space. NeuroImage, 162, 344–352. Retrieved from http://www.sciencedirect.com/science/article/pii/S1053811917306894 https://linkinghub.elsevier.com/retrieve/pii/S1053811917306894 doi:10.1016/j.neuroimage.2017.08.042

Billings, J., Thompson, G., Pan, W.-J., Magnuson, M. E., Medda, A., & Keilholz, S. (2018). Disentangling Multispectral Functional Connectivity With Wavelets. Frontiers in Neuroscience, 12, 812. Retrieved from https://www.frontiersin.org/article/10.3389/fnins.2018.00812/full doi: 10.3389/fnins.2018.00812

Biswal, B., Zerrin Yetkin, F., Haughton, V. M., & Hyde, J. S. (1995). Functional connectivity in the motor cortex of resting human brain using echo-planar mri. Magnetic Resonance in Medicine, 34(4), 537–541. Retrieved from https://doi.org/10.1002/mrm.1910340409 doi: 10.1002/mrm.1910340409

Buckner, R. L. K. F. M. C. A. D. J. C. Y. B. T. T. (2011). The organization of the human cerebellum estimated by intrinsic functional connectivity. Journal of Neurophysiology, 106(5), 2322–2345. Retrieved from http://jn.physiology.org/content/106/5/2322.abstract http://jn.physiology.org/content/jn/106/5/2322.full.pdf

Bullmore, E., & Sporns, O. (2009). Complex brain networks: graph theoretical analysis of structural and functional systems. Nat Rev Neurosci, 10(3), 186–198. doi: 10.1038/nrn2575

Calhoun, V. D., Miller, R., Pearlson, G., & Adalı, T. (2014). The chronnectome: time-varying connectivity networks as the next frontier in fmri data discovery. Neuron, 84(2), 262–274.

Carlsson, G. (2009). Topology and Data (Vol. 46; Tech. Rep. No. 2). Retrieved from http://www.ams.org/journals/bull/2009-46-02/S0273-0979-09-01249-X/S0273-0979-09-01249-X.pdf

Carrì, M., Cuturi, M., & Oudot, S. (n.d.). Sliced Wasserstein Kernel for Persistence Diagrams (Tech. Rep.). Retrieved from https://arxiv.org/pdf/1706.03358.pdf

Chaudhuri, R., Gerçek, B., Pandey, B., Peyrache, A., & Fiete, I. (2019). The intrinsic attractor manifold and population dynamics of a canonical cognitive circuit across waking and sleep. Nature Neuroscience, 22(9), 1512–1520. Retrieved from https://doi.org/10.1038/s41593-019-0460-x doi: 10.1038/s41593-019-0460-x

Chung, M. K., Lee, H., Di Christofano, A., Ombao, H., & Solo, V. (2019). Exact topological inference of the resting-state brain networks in twins. Network Neuroscience, 3(3), 674–694. Retrieved from https://www.mitpressjournals.org/doi/abs/10.1162/netn{_}a{_}00091 doi: 10.1162/netn_a_00091

Curto, C. (2017). What can topology tell us about the neural code? Bulletin of the American Mathematical Society, 54(1), 63–78. Retrieved from https://arxiv.org/pdf/1605.01905.pdf doi: 10.1090/bull/1554

Dabaghian, Y., Brandt, V. L., & Frank, L. M. (2014). Reconceiving the hippocampal map as a topological template. eLife, 3, e03476. doi: 10.7554/eLife.03476

Damoiseaux, J. S., & Greicius, M. D. (2009). Greater than the sum of its parts: a review of studies combining structural connectivity and resting-state functional connectivity. Brain Structure and Function, 213(6), 525–533.

De Silva, V., Morozov, D., & Vejdemo-Johansson, M. (2011). Dualities in persistent (co)homology (Vol. 27; Tech. Rep. No. 12). Retrieved from https://www.mrzv.org/publications/dualities-persistence/manuscript/ doi: 10.1088/0266-5611/27/12/124003

Duman, A. N., Tatar, A. E., Pirim, H., Duman, A. N., Tatar, A. E., & Pirim, H. (2019). Uncovering Dynamic Brain Reconfiguration in MEG Working Memory n-Back Task Using Topological Data Analysis. Brain Sciences, 9(6), 144. Retrieved from https://www.mdpi.com/2076-3425/9/6/144 doi: 10.3390/brainsci9060144

Edelsbrunner, H., Letscher, D., & Zomorodian, A. (2002). Topological persistence and simplification. Discrete and Computational Geometry, 28(4), 511–533. Retrieved from http://link.springer.com/10.1007/s00454-002-2885-2 doi: 10.1007/s00454-002-2885-2

Farahani, F. V., Karwowski, W., & Lighthall, N. R. (2019). Application of Graph Theory for Identifying Connectivity Patterns in Human Brain Networks: A Systematic Review. Frontiers in neuroscience, 13, 585. Retrieved from http://www.ncbi.nlm.nih.gov/pubmed/31249501 http://www.pubmedcentral.nih.gov/articlerender.fcgi?artid=PMC6582769 doi: 10.3389/fnins.2019.00585

Farge, M., & Marie. (1992). Wavelet Transforms and their Applications to Turbulence. Annual Review of Fluid Mechanics, 24(1), 395–458. Retrieved from http://www.annualreviews.org/doi/10.1146/annurev.fl.24.010192.002143 doi: 10.1146/annurev.fl.24.010192.002143

Finn, E. S., Shen, X., Scheinost, D., Rosenberg, M. D., Huang, J., Chun, M. M., … Constable, R. T. (2015). Functional connectome fingerprinting: identifying individuals using patterns of brain connectivity. Nature neuroscience, 18(11), 1664–1671.

Giusti, C., Ghrist, R., & Bassett, D. S. (2016). Two’s company, three (or more) is a simplex. Journal of Computational Neuroscience, 41(1), 1–14.

Giusti, C., Pastalkova, E., Curto, C., & Itskov, V. (2015). Clique topology reveals intrinsic geometric structure in neural correlations. Proceedings of the National Academy of Sciences, 112(44), 13455–13460. Retrieved from http://arxiv.org/abs/1502.06172{%}0A http://dx.doi.org/10.1073/pnas.1506407112 http://www.pnas.org/lookup/doi/10.1073/pnas.1506407112 doi: 10.1073/pnas.1506407112

Glasser, M. F., Coalson, T. S., Robinson, E. C., Hacker, C. D., Harwell, J., Yacoub, E., … Van Essen, D. C. (2016). A multi-modal parcellation of human cerebral cortex. Nature, 536(7615), 171–178. Retrieved from http://www.ncbi.nlm.nih.gov/pubmed/27437579 http://www.pubmedcentral.nih.gov/articlerender.fcgi?artid=PMC4990127 http://www.nature.com/articles/nature18933 doi: 10.1038/nature18933

Gonzalez-Castillo, J., Hoy, C. W., Handwerker, D. A., Robinson, M. E., Buchanan, L. C., Saad, Z. S., & Bandettini, P. A. (2015). Tracking ongoing cognition in individuals using brief, whole-brain functional connectivity patterns. Proceedings of the National Academy of Sciences of the United States of America, 112(28), 8762–7. Retrieved from http://www.ncbi.nlm.nih.gov/pubmed/26124112 http://www.pubmedcentral.nih.gov/articlerender.fcgi?artid=PMC4507216 doi: 10.1073/pnas.1501242112

Gordon, E. M., Laumann, T. O., Adeyemo, B., Huckins, J. F., Kelley, W. M., & Petersen, S. E. (2016). Generation and Evaluation of a Cortical Area Parcellation from Resting-State Correlations. Cerebral cortex (New York, N.Y.: 1991), 26(1), 288–303. Retrieved from http://www.ncbi.nlm.nih.gov/pubmed/25316338 http://www.pubmedcentral.nih.gov/articlerender.fcgi?artid=PMC4677978 doi: 10.1093/cercor/bhu239

Guerra, M., De Gregorio, A., Fugacci, U., Petri, G., & Vaccarino, F. (2020). Homological scaffold via minimal homology bases. arXiv preprint arXiv:2004.11606.

Hansen, E. C., Battaglia, D., Spiegler, A., Deco, G., & Jirsa, V. K. (2015). Functional connectivity dynamics: modeling the switching behavior of the resting state. Neuroimage, 105, 525–535.

Hutchison, R. M., Womelsdorf, T., Allen, E. A., Bandettini, P. A., Calhoun, V. D., Corbetta, M., … others (2013). Dynamic functional connectivity: promise, issues, and interpretations. Neuroimage, 80, 360–378.

Ibáñez-Marcelo, E., Campioni, L., Phinyomark, A., Petri, G., & Santarcangelo, E. L. (2019). Topology highlights mesoscopic functional equivalence between imagery and perception: The case of hypnotizability. NeuroImage, 200, 437–449. Retrieved from https://www.sciencedirect.com/science/article/pii/S1053811919305415?via{%}3Dihub doi: 10.1016/J.NEUROIMAGE.2019.06.044

Kunert-Graf, J. M., Eschenburg, K. M., Galas, D. J., Kutz, J. N., Rane, S. D., & Brunton, B. W. (2019). Extracting Reproducible Time-Resolved Resting State Networks Using Dynamic Mode Decomposition. Frontiers in Computational Neuroscience, 13, 75. Retrieved from https://www.frontiersin.org/article/10.3389/fncom.2019.00075/full doi: 10.3389/fncom.2019.00075

Lerda, G. (2016). University of Torino, Torino, Italy (Unpublished doctoral dissertation). University of Turino.

Lord, L. D., Expert, P., Fernandes, H. M., Petri, G., Van Hartevelt, T. J., Vaccarino, F., … Kringelbach, M. L. (2016). Insights into brain architectures from the homological scaffolds of functional connectivity networks. Frontiers in Systems Neuroscience, 10(NOV). doi: 10.3389/fnsys.2016.00085

Lurie, D. J., Kessler, D., Bassett, D. S., Betzel, R. F., Breakspear, M., Kheilholz, S., … Calhoun, V. D. (2020). Questions and controversies in the study of time-varying functional connectivity in resting fMRI. Network Neuroscience, 4(1), 30–69. Retrieved from https://www.mitpressjournals.org/doi/abs/10.1162/netn{_}a{_}00116 doi: 10.1162/netn_a_00116

Marwan, N., Thiel, M., & Nowaczyk, N. R. (2002). Application of the cross wavelet transform and wavelet coherence togeophysical time series. Nonlinear Processes in Geophysics, 9(3), 325–331. Retrieved from http://sub3.isiknowledge.com/error/Error?PathInfo=/&Domain=isiknowledge.com&Params=DestApp=WOS&DestParams={%}3Faction{%}3Dretrieve{%}26mode{%}3DFullRecord{%}26product{%}3DWOS{%}26UT{%}3D0001769 http{%}3A{%}2F{%}2Faccess.isiproducts.co doi: 10.5194/npg-11-515-2004

McInnes, L., Healy, J., & Melville, J. (2020). Umap: Uniform manifold approximation and projection for dimension reduction.

Mileyko, Y., Mukherjee, S., & Harer, J. (2011). Probability measures on the space of persistence diagrams. Inverse Problems, 27(12), 124007. Retrieved from https://iopscience.iop.org/article/10.1088/0266-5611/27/12/124007 doi:10.1088/0266-5611/27/12/124007

Patania, A., Vaccarino, F., & Petri, G. (2017). Topological analysis of data. EPJ Data Science, 6, 1–6.

Petri, G., Expert, P., Turkheimer, F., Carhart-Harris, R., Nutt, D., Hellyer, P. J., & Vaccarino, F. (2014a). Homological scaffolds of brain functional networks. Journal of The Royal Society Interface, 11(101), 20140873.

Petri, G., Expert, P., Turkheimer, F., Carhart-Harris, R., Nutt, D., Hellyer, P. J., & Vaccarino, F. (2014b). Homological scaffolds of brain functional networks. Journal of The Royal Society Interface, 11(101), 20140873–20140873. Retrieved from http://dx.doi.org/10.1098/rsif.2014.0873orvia http://rsif.royalsocietypublishing.org. http://rsif.royalsocietypublishing.org/cgi/doi/10.1098/rsif.2014.0873 doi:10.1098/rsif.2014.0873

Petri, G., Scolamiero, M., Donato, I., & Vaccarino, F. (2013). Topological strata of weighted complex networks. PloS one, 8(6).

Phinyomark, A., Ibanez-Marcelo, E., & Petri, G. (2017). Resting-state fmri functional connectivity: Big data preprocessing pipelines and topological data analysis. IEEE Transactions on Big Data, 3(4), 415–428.

Rasetti, M. (2017). The ‘Life Machine’: A Quantum Metaphor for Living Matter. International Journal of Theoretical Physics, 56(1), 145–167. doi: 10.1007/s10773-016-3177-6

Reimann, M. W., Nolte, M., Scolamiero, M., Turner, K., Perin, R., Chindemi, G., … Markram, H. (2017). Cliques of Neurons Bound into Cavities Provide a Missing Link between Structure and Function. Frontiers in Computational Neuroscience, 11, 48. Retrieved from http://journal.frontiersin.org/article/10.3389/fncom.2017.00048/full https://www.frontiersin.org/articles/10.3389/fncom.2017.00048/full doi: 10.3389/fncom.2017.00048

Saggar, M., Sporns, O., Gonzalez-Castillo, J., Bandettini, P. A., Carlsson, G., Glover, G., & Reiss, A. L. (2018). Towards a new approach to reveal dynamical organization of the brain using topological data analysis. Nature Communications, 9(1). Retrieved from www.nature.com/naturecommunications doi: 10.1038/s41467-018-03664-4

Sizemore, A., Giusti, C., & Bassett, D. S. (2017). Classification of weighted networks through mesoscale homological features. Journal of Complex Networks, 5(2), 245–273.

Sizemore, A. E., Giusti, C., Kahn, A., Vettel, J. M., Betzel, R. F., & Bassett, D. S. (2018). Cliques and cavities in the human connectome. Journal of Computational Neuroscience, 44(1), 115–145. Retrieved from https://arxiv.org/pdf/1608.03520.pdf doi: 10.1007/s10827-017-0672-6

Sizemore, A. E., Phillips-Cremins, J. E., Ghrist, R., & Bassett, D. S. (2019). The importance of the whole: Topological data analysis for the network neuroscientist. Network Neuroscience, 3(3), 656–673. Retrieved from https://doi.org/10.1162/netn{_}a{_}00073 doi: 10.1162/netn_a_00073

Smith, S. M., Fox, P. T., Miller, K. L., Glahn, D. C., Fox, P. M., Mackay, C. E., … Beckmann, C. F. (2009). Correspondence of the brain’s functional architecture during activation and rest. Proceedings of the National Academy of Sciences. Retrieved from http://www.pnas.org/content/early/2009/07/17/0905267106.abstract http://www.pnas.org/content/106/31/13040.full.pdf doi: 10.1073/pnas.0905267106

Stolz, B. (2014). Computational Topology in Neuroscience (Doctoral dissertation). Retrieved from https://pdfs.semanticscholar.org/62b0/f4a74b73c9dd3948a63189d2b3f3cfc7185d.pdf

Stoodley, C. J., Valera, E. M., & Schmahmann, J. D. (2010). An fMRI study of intra-individual functional topography in the human cerebellum. Behavioural Neurology, 23(1-2), 65–79. Retrieved from https://pubmed.ncbi.nlm.nih.gov/20714062/ doi: 10.3233/BEN-2010-0268

Thompson, W. H., & Fransson, P. (2015). The frequency dimension of fMRI dynamic connectivity: Network connectivity, functional hubs and integration in the resting brain. Neuroimage, 121, 227–242. doi: 10.1016/j.neuroimage.2015.07.022

Torrence, C., Compo, G. P., Torrence, C., & Compo, G. P. (1998). A Practical Guide to Wavelet Analysis. Bulletin of the American Meteorological Society, 79(1), 61–78. Retrieved from http://journals.ametsoc.org/doi/abs/10.1175/1520-0477{%}281998{%}29079{%}3C0061{%}3AAPGTWA{%}3E2.0.CO{%}3B2 doi: 10.1175/1520-0477(1998)0790061:APGTWA2.0.CO;2

Torrence, C., Webster, P. J., Torrence, C., & Webster, P. J. (1999). Interdecadal Changes in the ENSO–Monsoon System. Journal of Climate, 12(8), 2679–2690. Retrieved from http://journals.ametsoc.org/doi/abs/10.1175/1520-0442{%}281999{%}29012{%}3C2679{%}3AICITEM{%}3E2.0.CO{%}3B2 doi:10.1175/1520-0442(1999)0122679:ICITEM2.0.CO;2

Tralie, C., Saul, N., & Bar-On, R. (2018). Ripser.py: A lean persistent homology library for python. The Journal of Open Source Software, 3(29), 925. Retrieved from https://doi.org/10.21105/joss.00925 doi: 10.21105/joss.00925

Yoo, J., Kim, E. Y., Ahn, Y. M., & Ye, J. C. (2016). Topological persistence vineyard for dynamic functional brain connectivity during resting and gaming stages. Journal of Neuroscience Methods, 267, 1–13. doi: 10.1016/j.jneumeth.2016.04.001

